# Anatomical and functional connectivity support the existence of a salience network node within the caudal ventrolateral prefrontal cortex

**DOI:** 10.1101/2021.10.01.462813

**Authors:** Lucas R. Trambaiolli, Xiaolong Peng, Julia F. Lehman, Hesheng Liu, Suzanne N. Haber

**Affiliations:** McLean Hospital, Harvard Medical School, United States; University of Rochester School of Medicine & Dentistry, United States; Massachusetts General Hospital, Harvard Medical School, United States; Medical University of South Carolina, United States

**Author notes:** Address correspondence to: Dr. Suzanne Haber. Contributed equally to this manuscript (Lucas R. Trambaiolli, Xiaolong Peng). Contributed equally to this manuscript (Hesheng Liu, Suzanne N. Haber).

**Keywords:** attention, salience network, ventrolateral prefrontal cortex, tract-tracing, non-human primates, functional connectivity

## Abstract

Three large-scale brain networks are considered essential to cognitive flexibility: the ventral and dorsal attention (VAN and DAN) and salience (SN) networks. The ventrolateral prefrontal cortex (vlPFC) is a known component of the VAN and DAN, but there is an important gap in the current knowledge regarding its involvement in the SN. In this study, we used a translational and multimodal approach to fulfill this gap and demonstrate the existence of a SN node within the vlPFC. First, we used tract-tracing methods in non-human primates (NHP) to quantify the anatomic connectivity strength between the different vlPFC areas and the frontal and insular cortices. The strongest connections with the dorsal anterior cingulate cortex (dACC) and anterior insula (AI) locations comprising the two main cortical SN nodes were derived from the caudal area 47/12. This location also has strong axonal projections to subcortical structures of the salience network, including the dorsomedial thalamus, hypothalamus, sublenticular extended amygdala, and periaqueductal gray. Second, we used a seed-based functional connectivity analysis in NHP resting-state functional MRI (rsfMRI) data to validate the caudal area 47/12 as an SN node. Third, we used the same approach in human rsfMRI data to identify a homologous structure in caudal area 47/12, also showing strong connections with the SN cortical nodes, thus confirming the caudal area 47/12 as the SN node in the vlPFC. Taken together, the vlPFC contains nodes for all three cognitive networks, the VAN, DAN, and SN. Thus, the vlPFC is in a position to switch between these three cognitive networks, pointing to a key role as an attentional hub. Its tight additional connections to the orbitofrontal, dorsolateral, and ventral premotor cortices, places the vlPFC at the center for switching behaviors based on environmental stimuli, computing value and cognitive control.

## Introduction

Three distributed attentional networks, the dorsal and ventral attention (DAN and VAN) and salience (SN) networks, play key roles in switching actions based on environmental stimuli (1-3). The DAN is a top-down bilateral fronto-parietal network, responsible for *selecting* stimuli and responses (2, 4). The VAN is a bottom-up ventral fronto-parieto-temporal network, responsible for *detecting* outstanding stimuli and reorienting ongoing activity (2, 4). The salience network (SN) (3, 5), cortically anchored in the anterior insula (AI) and the dorsal anterior cingulate cortex (dACC), adds value to external and internal stimuli, driving attention to rapidly modify behaviors (6). The SN works closely with the VAN, to ‘pull’ attention to valued stimuli, based on a combination of previous experience and motivation. However, all three networks must operate together for rapid environmental responses. The ventrolateral prefrontal cortex (vlPFC) lies at the junction between the DAN (areas 44 and 45) (7-13) and VAN (area 47/12) (12-16). In contrast, based on imaging studies, the key nodes of the SN are ACC and AI, and not the vlPFC. Yet, the vlPFC, particularly area 47/12, is central for assessing value and, along with the ACC drives information seeking, to provide value-related discriminations (17). Indeed, it is the orbital portion of area 47/12 is involved in stimulus-outcome predictions (18-20), and, when lesioned, interferes with object reversal learning. Area 47/12 is tightly connected to both the ACC and the adjacent AI (21). However, area 47/12 is large and connected to a wide range of cortical regions. We posit that embedded within this large area is a separate SN node that links the ACC and AI with the vlPFC that has not been evident due to the technical limitation of functional MRI (3, 6, 22). We demonstrate here, that, based on its anatomic organization and connections to the two central nodes of the SN (dACC and AI) the vlPFC is a distinct node in the SN, separate from the adjacent AI. We also show that, with anatomic guidance, this separate node can be identified using fMRI in the human brain. A SN component within the vlPFC brings unique information about stimulus value to this network, through its connections with the orbitofrontal cortex and thus complementary to the roles of the AI and dACC in information integration and information seeking, respectively. Given the high interconnectivity of areas 44, 45, and 47/12, a SN node within the vlPFC places it in a central hub-like position to integrate information across the three main attention networks, supporting the region’s central role in modulating behavioral flexibility (23-25).

We used a cross-species and cross-modality approach to determine the relative strengths of connections of subregions of the vlPFC with the two SN cortical nodes, the AI and ACC, compared to other frontal regions: tract-tracing methods in macaque monkeys, followed by a seed-based fMRI approach to determine connectivity strength first in the NHP then in humans. We first quantified the anatomic connectivity strength between the different vlPFC subregions and the frontal and insular cortices. We found that the strongest connections with the dACC and AI were with the caudal area 47/12. This sublocation also presented strong axonal projections to subcortical structures of the salience network, including the dorsomedial thalamus (DT), sublenticular extended amygdala (SEA), substantia nigra/ventral tegmental area (SN/VTA), and periaqueductal gray (PAG). Using resting-state functional connectivity MRI (fcMRI), we found that the connectivity strength and patterns between the subregions of the vlPFC and the dACC and AI SN nodes were similar to anatomic data in NHP. Finally, placing seeds in homologous vlPFC regions in the human, we show that, similar to the NHP results, fcMRI connectivity between caudal 47/12 is significantly stronger with the dACC and AI compared to other vlPFC regions.

## Results

### Retrograde tracing reveals a SN node in the caudal area 47/12

Retrograde tracing injections were placed in areas 44, 45 and subregions of 47/12 on the right vlPFC (coronal representations of injection centers and 3D view of injections in Supp. Fig. 1) and the labeled cells in the frontal and insular cortices were charted. We focused on the right hemisphere to reduce the effect of species specificities associated to language development in our analyses (26). To determine the relative projection strengths across cases, we calculated the percentage of total labelled cells that project from each cytoarchitectonic area to each injection site. To compare the projection strengths to what would be expected by chance, we performed a random sampling analysis by permuting neurons 10^6^ times among each frontal or insular cortex area with a probability given by the volume of each area. To evaluate the strength of connections from the main cortical nodes of the SN, we compared projections from the dACC (area 24) and AI (areas OPAl, OPro, IPro and AI) across cases. The results demonstrate that the connectivity strength varies across vlPFC areas (Fig. 1A, extended bar charts are shown in Supp. Fig. 2).

**Fig. 1.**
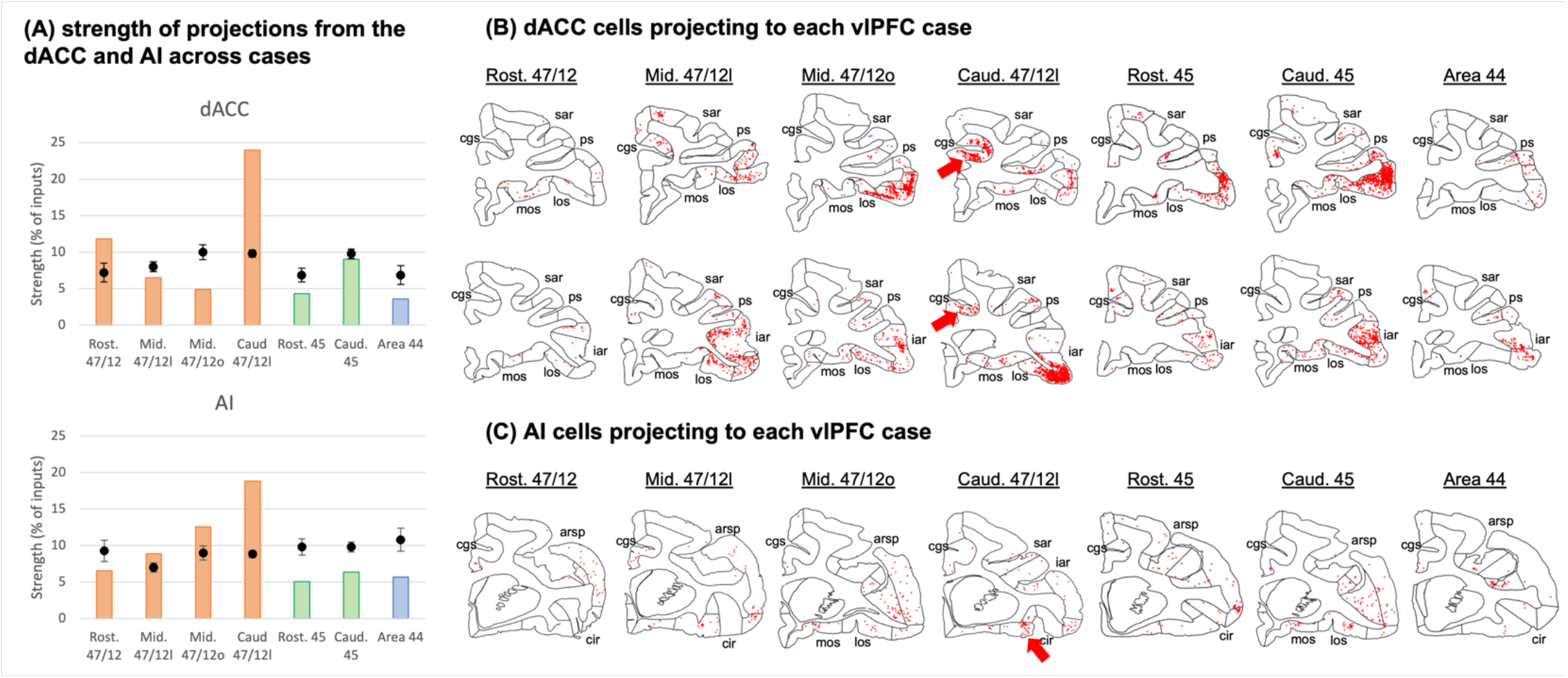
Strength of projections from Salience Network cortical nodes grouped by cytoarchitectonic divisions across cases. **(A)** The dACC correspond to area 24, while the AI is the combination of areas OPAl, Opro, IPro and AI. Orange bars illustrate cases with injections in area 47/12, green bars in area 45, and blue bars in area 44. Black dots show the average and standard-deviation of random sampling from the respective areas in each case. Coronal sections and the respective labelled cells (red dots) in the **(B)** dACC and **(C)** AI projecting to the caudal area 47/12 in the vlPFC. The black circles represent the areas of interest for the Salience Network. *Abbreviations:* arsp = arcuate sulcus spur; cgs = cingulate sulcus; cir = circular sulcus; iar = inferior arcuate sulcus; los = lateral orbital sulcus; mos = medial orbital sulcus; ps = principal sulcus; sar = superior arcuate sulcus.

**Fig. 2.**
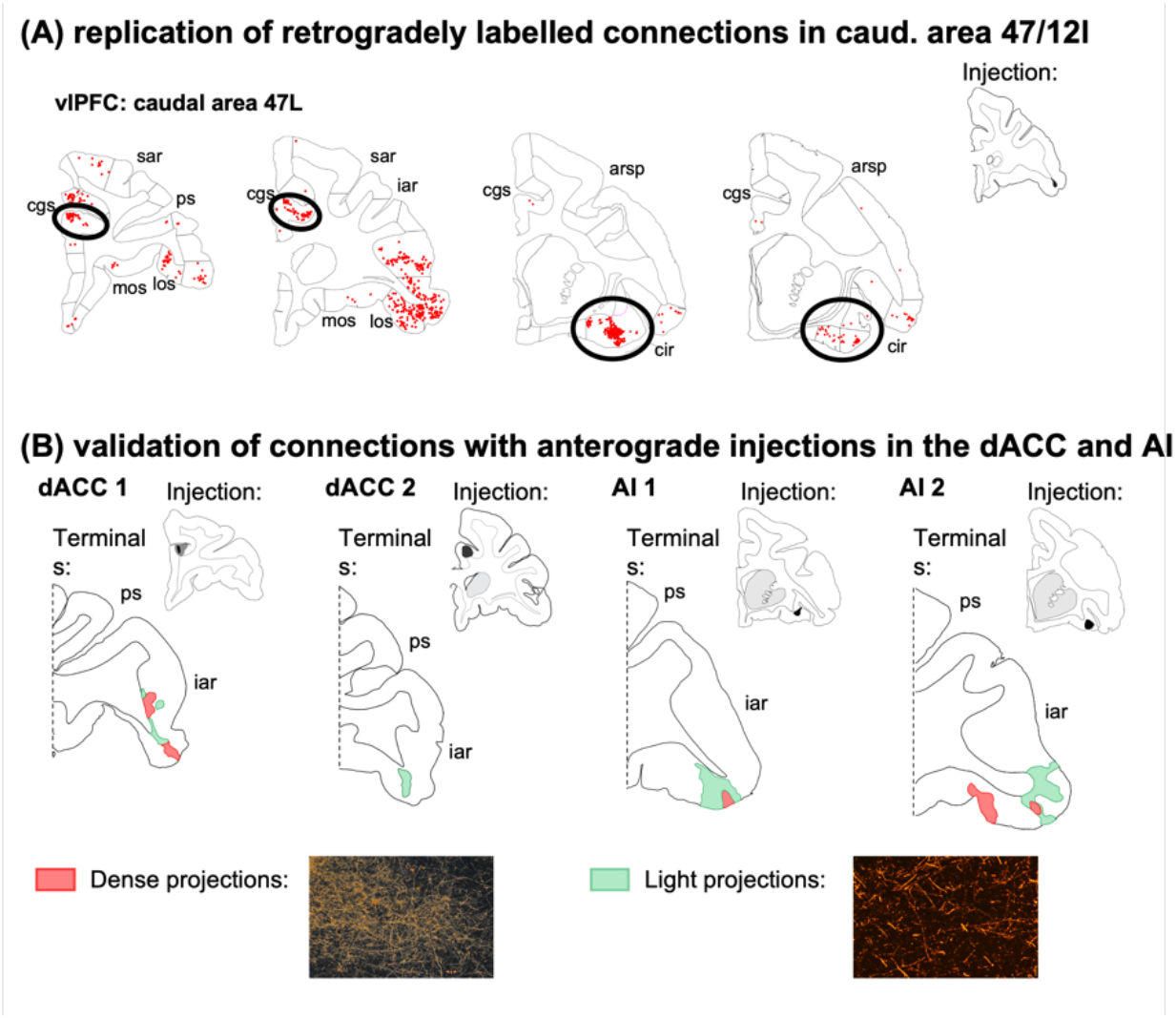
Anatomical replication and validation of the caudal vlPFC as a salience network node. **(A)** Coronal sections and the respective labelled cells (red dots) from the validation vlPFC retrograde tracing injection (case 4b). The connectivity strength and correlation between both caudal vlPFC cases is shown in the bar graph (orange bars correspond to case 4a, and blue bars to case 4b). **(B)** Coronal sections and the respective labelled terminal fields from the validation anterograde tracer injections in the dACC and AI (red areas correspond to dense axonal projections and green areas to light axonal projections). *Abbreviations:* arsp = arcuate sulcus spur; cgs = cingulate sulcus; cir = circular sulcus; iar = inferior arcuate sulcus; los = lateral orbital sulcus; mos = medial orbital sulcus; ps = principal sulcus; sar = superior arcuate sulcus.

Among all vlPFC injections, caudal area 47/12l stands out as the main location for connections from the dACC and the AI. This area, in addition to rostral 47/12, showed connectivity strength above the chance level with dACC (area 24). Specifically, area 24 projections to caudal area 47/12 were at least twice as strong as expected by chance and twice as strong compared to the projections to the other vlPFC locations. Clusters of projecting cells were found in both pre- and post-genual dACC (Fig. 1B). For projections from the AI, caudal area 47/12l had the highest difference from the chance level, twice as high compared with injections in mid 47/12. Interestingly, these cells clusters are located in the orbital portion around the beginning of the circular sulcus in the AI. Specifically, this region in the macaque brain is enriched with von Economo neurons (27) (Fig. 1C), a cell type rare in the brain but characteristic of the SN (3, 6). These data demonstrate that a specific vlPFC region, caudal 47/12, is tightly linked to the two SN nodes. Projection patterns from other cortical areas are shown in Supp. Fig. 3.

**Fig. 3.**
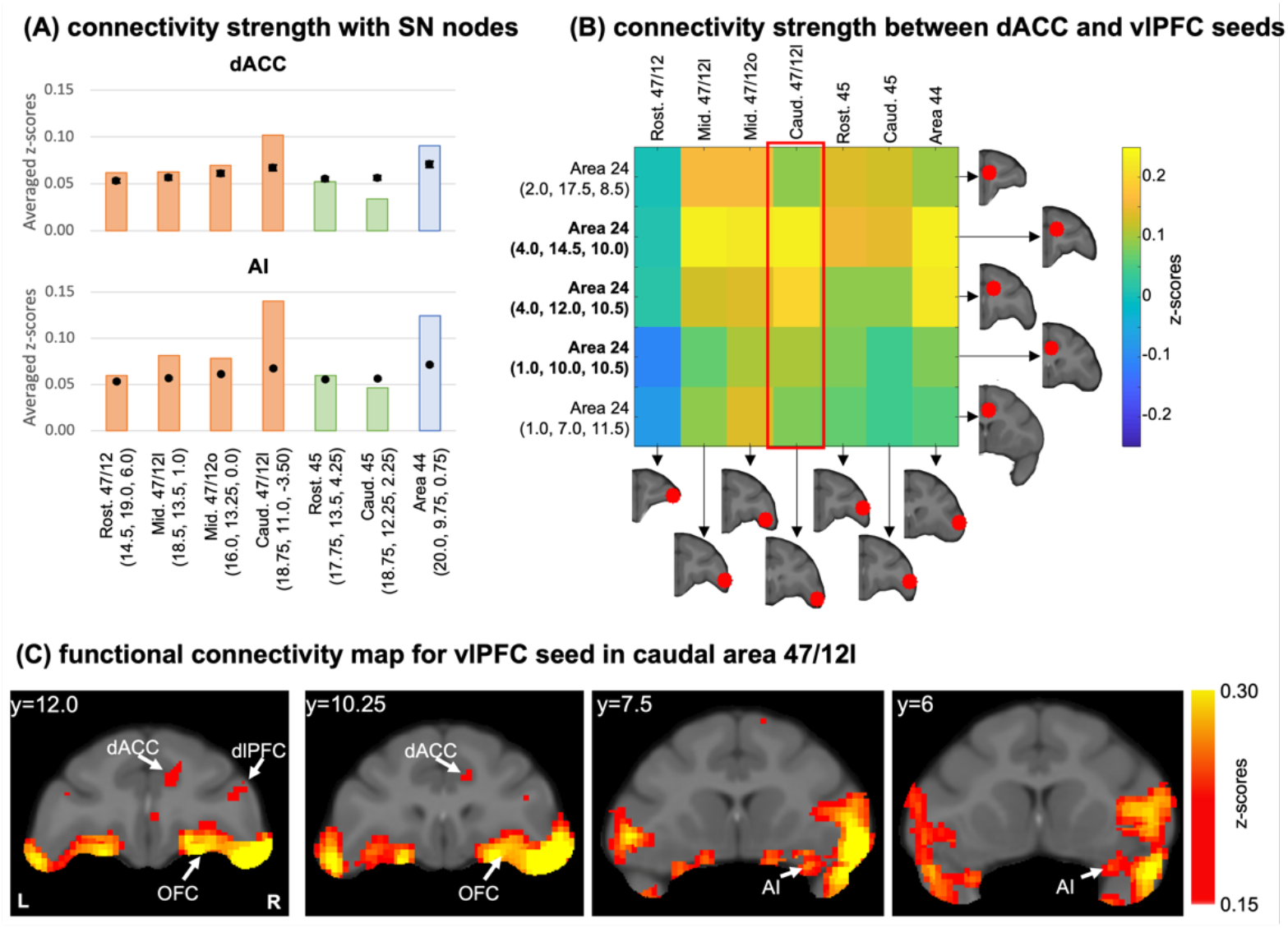
Functional connectivity analysis in the macaque brain. **(A)** Average connectivity strength (z-scores) between vlPFC seeds and the dACC and AI masks. Orange bars illustrate cases with injections in area 47/12, green bars in area 45, and blue bars in area 44. Black dots show the average and standard-deviation of the voxel permutation analysis. **(B)** Connectivity strength (z-scores) between dACC and vlPFC seeds. In bold the seeds overlapping with the dACC mask. **(C)** Different views of the voxel distribution for the caudal 47/12l seed.

To replicate these results, we placed an additional retrograde injection at a similar vlPFC location and found clusters of labeled cells in the same positions within the dACC and AI (Fig. 2A). Moreover, this injection site was highly correlated with the original caudal 47/12 injection in regards of overall distribution of connectivity strengths across the frontal and insular cortices (rho = 0.70, p << 0.01). To verify the convergence of dACC and AI inputs to the caudal area 47/12l, small anterograde tracer injections were placed at same location as the clusters of dACC and AI labeled cells (Fig. 2B). Fibers from these injection sites terminated in the caudal area 47/12l. These results are consistent with similar injections within the vlPFC, dACC and AI reported in qualitative studies (21, 28-32), and support our findings that there are convergent inputs from the dACC and AI to specific regions of the vlPFC.

The SN is also characterized by specific subcortical connections, including the SEA, ventral striatum (VS), DT, hypothalamus, SN/VTA, and PAG (3, 5, 6). Importantly, following anterograde injections into caudal area 47/12l. Area 44 has light terminal labeling in DT, hypothalamus, and SN/VTA, but not in the SEA and VS. In area 45 terminals were predominantly found in DT, but not in other subcortical nodes. Rostral and mid 47/12 have terminals in DT and SEA. Mid 47/12 also lightly projected to the SN/VTA and lateral hypothalamus. Caudal 47/12 had a particular combination of projections, with dense terminal fields located in the SEA, DT, SN/VTA, hypothalamus, and PAG (Supp. Fig. 4). There were fibers and terminals located along the base of the brain streaming through the SEA, with some terminating in the lateral hypothalamus. Moreover, dense terminals fields were also located in the DT, with fewer fibers in the PAG. However, consistent with previous cortico-striatal studies, there were no fibers in the VS. Indeed, vlPFC fibers terminate dorsal to the VS stretching from the ventral rostral putamen and to the central caudate nucleus, just dorsal to the VS (33-35). These connections are consistent with previous anatomical studies (36-38), and provide additional evidence endorsing the role of the caudal area 47/12l in the SN.

**Fig. 4.**
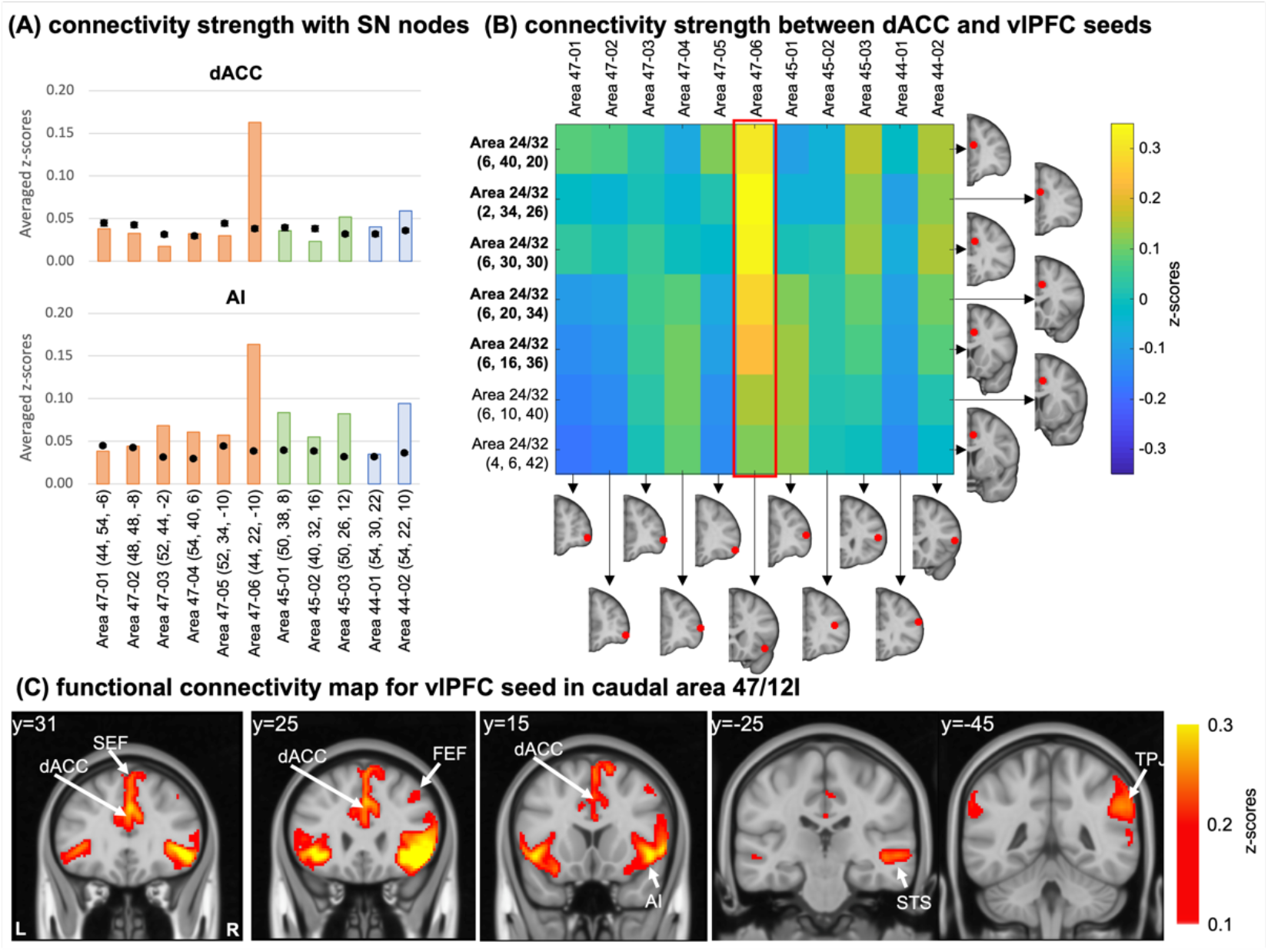
Functional connectivity analysis in the human brain. **(A)** Average connectivity strength (z-scores) between vlPFC seeds and the dACC and AI masks. Orange bars illustrate cases with injections in area 47/12, green bars in area 45, and blue bars in area 44. Black dots show the average and standard-deviation of the voxel permutation analysis. **(B)** Connectivity strength (z-scores) between dACC and vlPFC seeds. In bold the seeds overlapping with the dACC mask. **(C)** Different views of the voxel distribution for the caudal 47/12l seed. All coordinates are in the human MNI space.

### The SN node within the caudal vlPFC can be identified using NHP fcMRI

We then investigated how well these anatomical connectivity patterns may correspond to resting state functional connectivity patterns measured by fMRI. Using data from the PRIMatE Data Exchange (PRIME-DE) consortium (39), we placed seven seeds of 3 mm radius in matched locations to our anatomic injection sites and calculated the functional connectivity between each seed and all brain voxels. Masks for the dACC and AI were created with reference to the clusters of cells observed in the retrograde data. The connectivity strength was computed as the average of absolute connectivity values inside each mask. We also performed 10^6^ random permutations of voxels across the brain volume and computed the random distribution of connectivity strengths in each mask.

The functional connectivity pattern between each vlPFC seed and the dACC mask (Fig. 3A, top) showed strength around or below the chance level in rostral and mid area 47/12 and area 45. Connectivity strength in caudal area 47/12 and area 44 were above the chance and stands out compared to other brain regions. For functional connectivity between the vlPFC seeds and the AI mask (Fig. 3A, bottom), a rostro-caudal gradient is observed in area 47/12, with the caudal area 47/12 presenting the highest connectivity strength among all locations. Again, area 45 showed connection strength below the chance level, while area 44 was above the chance.

To ensure the strong connections with the dACC are not artifacts given the proximity of the vlPFC seeds to the AI, we performed a complementary analysis placing 5 seeds within the right dACC (inside and outside the mask). Then, we calculated the seed-to-seed functional connectivity between the dACC and vlPFC (Fig 3B). Consistent with the mask analysis, the caudal area 47/12 and area 44 showed the strongest connections with the dACC seeds within the mask. These connectivity profiles are overall consistent with the anatomical data, with exception to area 44, which did not show strong connections based on the anatomic tracing (see Fig. 1A). The results from the fcMRI in area 44 are likely due to the proximity with the caudal area 47/12 and overlap between these seeds. Fig. 3C shows the location of voxels within the dACC and AI with high functional connectivity with the seed in caudal area 47/12.

### A vlPFC salience network node in the human functional connectivity map

To translate the results from NHP fcMRI to human fcMRI analysis we placed eleven seeds of 5 mm radius across the vlPFC areas. We calculated the functional connectivity between each seed and all brain voxels from 1000 healthy adult subjects from a publicly available, fully preprocessed dataset (Brain Genomics Superstruct Project) (40). Masks for the dACC and AI were created outlining regions homologous to those containing clusters of cells (Fig. 1B-C; Fig. 2A) (41). The computation of connectivity strength and voxel permutation analysis followed the same approach used for the monkey data.

Overall connectivity strength with the dACC mask (Fig. 4A, top) was below, or around, the chance level, with exception of the caudal-most seeds in each vlPFC area. Specifically, the strongest connection was with the caudal area 47, similar to the results in the NHP anatomy and imaging data. The connectivity pattern observed between each seed and the AI mask (Fig. 4A, bottom) is also consistent with the patterns observed in the NHP anatomy. Specifically, connectivity strengths between AI and area 47 are organized in a light rostro-caudal gradient, with the strongest connection in caudal area 47. This gradient is also consistent with the pattern observed in the NHP tracing (Fig 1A) and fcMRI data (Fig 4A, bottom). Areas 45 and 44 also presented connectivity strengths with the AI and dACC masks around the chance level for the dACC mask and slightly higher for the AI. For all these cases, however, the observed strengths were still lower than the caudal area 47/12l. Additionally, we placed 7 seeds within the right dACC (inside and outside the mask) and calculated the seed-to-seed functional connectivity between dACC and vlPFC seeds (Fig 4B). As expected, the vlPFC seed in caudal area 47 was the one showing strongest connectivity with the dACC seeds within and around the created mask. These results are consistent with the NHP anatomical and fcMRI data, support the caudal area 47/12 as a node of the SN. The translation of these nodes and connections across species is supported by anatomic-functional homologies within the vlPFC of humans and NHP (21, 42). We repeated the experiment using 3 mm seeds to show that these results are independent to the seed size and possible overlapping of the original seed with beginning of the insular cortex (Supp. Fig. 5). Finally, we found high functional connectivity between the caud 47/12l seed and a cluster of voxels in the dACC and AI, two main cortical nodes of the SN (3, 6) (Fig. 4C).

## Discussion

### Summary

The presence of salient stimuli that activates the SN also activates the vlPFC (43-47). However, due to inherent limitations of functional MRI in deciphering signal locations between adjacent cortical areas, the vlPFC component of the SN is largely ignored, with the assumption that activation is simply part of the AI signal (3, 6, 22). This assumption has had important ramifications for understanding, not only the SN, but also how the three attention networks might be anatomically linked. In this study, we provide cross-modal and cross-species evidence, based on connectivity, for a separate SN node located in the caudal area 47/12 within the right vlPFC. This region showed a peak of anatomical and functional connectivity with the main cortical nodes (dACC and AI), as well as anatomical projection to subcortical nodes of the SN (DT, hypothalamus, SN/VTA, SEA, and PAG). In addition to extending our understanding on the structure of the SN, our experiment also provides an important methodological contribution to mapping large-scale brain networks. Although fMRI is useful to provide a general view of these circuits, only the precision of NHP tracing is capable to describe the specificities of individual connections, and how they are characterized in each network (48), as demonstrated here.

### Caudal vlPFC is a node in the SN

The proposed inclusion of caudal 47/12 in the SN is primarily based on two lines of anatomic evidence: first, the presence of direct monosynaptic connections to specific regions within the two main cortical SN nodes, the dACC and AI; and second, a pattern of connections with subcortical areas that are also considered part of the SN. At the cortico-cortical level, our innovative combination of neuroanatomical tracing methods in NHP with random sampling analysis showed that this area is tightly linked to the dACC and AI. We identified anatomical connectivity strengths significantly above chance levels for each vlPFC subregion and calculated the strength of inputs from the two main cortical nodes of the SN (dACC and AI). Importantly, we had several injections in area 47/12, which is a particularly large region that can be further subdivided based on connectivity (21, 49-51). The peak of connections from both the dACC and AI to the vlPFC specifically targeted the caudal 47/12. In fact, the strength of the dACC connections was twice as high as connections to other vlPFC subdivisions. Anterograde injections in the dACC corroborated the existence and strength of these connections to caudal 47/12. The cluster of cells from AI projecting to caudal 47/12 was identified predominantly in the rostral portions of the AI. Anterograde injections in this rostral AI region confirmed its connections with caudal 47/12. This AI region is also characterized by the presence of a group of unique neurons (von Economo neurons - VENs), in both humans and NHPs (27, 52). VENs have distinctive properties, including fast axonal electric conduction between projected areas (53), which allows for quick identification of salient stimuli (3, 6). Importantly, VENs are predominantly found in the right hemisphere compared to the left (27, 52), the same hemisphere of the caudal 47/12 SN node candidate.

The SN also has subcortical components: the DT, hypothalamus, SN/VTA, SEA, VS, and PAG (3, 5). We found that most sections of vlPFC displayed partial connectivity to subcortical nodes. Specifically, all areas projected axon terminals to DT, but connections with other subcortical regions varied per vlPFC location. Area 44 projected to the DT, lateral hypothalamus and SN/VTA, but not in the SEA and VS. Area 45 terminals were predominantly found in DT, but not in other subcortical nodes. Mid and caudal 47/12 projected to the DT, lateral hypothalamus, SEA, and SN/VTA, but not to VS. However, caudal 47/12 projections were denser than those observed from mid 47/12. Caudal 47/12 stands out from mid 47/12 given its combination of strong connections with the cortical SN nodes, and dense projections to the subcortical nodes, providing further support that this location is part of the SN.

We further translated these tracing results by probing their consistency with fcMRI in NHP (48). The seed placement in the NHP corresponded to the injection locations. We computed the connectivity strength with two cortical masks created corresponding to the cell clusters in the AI and dACC. As expected, caudal 47/12 showed the highest connectivity strength with both the dACC and AI SN nodes. These results were replicated when placing seeds within the dACC. However, one potential limitation of our analysis is the existence of a peak of rsFC between the area 44 and the cortical SN nodes. A possible reason for this result is the spatial overlapping between seeds in caudal 47/12 and area 44, given the resolution of the MRI data available. This resolution limitation highlights the advantages of cross-modality comparisons within the same species when finely delineating brain connectivity to avoid misleading conclusions (48). Similar patterns of vlPFC connectivity were observed when we systematically placed seeds throughout the human vlPFC. Seeds placed in caudal area 47/12 showed the maximum connectivity strength with both dACC and AI masks, consistent with our anatomical and imaging results in NHP. When placing seeds within the dACC mask for a seed-to-seed analysis, caudal 47/12 again showed the strongest connections with dACC subareas. One important aspect of our results is that, in both NHP and humans, caudal 47/12 connections to subcortical SN nodes were not as distinguishable as in the tracing data. This limitation is somehow expected. A previous study using a seed-based rsFC approach to replicate large-scale networks also reported weaker subcortical connections within the SN (13).

This cross-modality and cross-species study provides empirical evidence that caudal area 47/12 is anatomically and functionally connected with the SN. This location in humans is within the vlPFC area mistakenly merged with the AI into the fronto-insular cortex (FIC) definition (3, 22). However, caudal area 47/12 and AI are separate structural entities, with different anatomical organization and connectivity profiles (21, 31, 32, 54-56). Together, these data support that caudal 47/12 should be considered as an independent SN node, separate from the original AI/FIC definition.

### Possible roles of caudal vlPFC within the SN

In addition to identifying salient stimuli, the SN recruits behaviorally appropriate responses (3, 57). For this purpose, each cortical SN node has a specific function. The dACC is related to action selection (58, 59), given its connections with motor control regions (30). The AI is the node combining sensorial, interoceptive, and limbic information to process salient stimuli (59, 60) due to its cortico-cortical connections with sensory and limbic regions (61, 62). In addition to projections from AI and dACC, caudal area 47/12 is connected with sensory areas in the temporal pole, cognitive control regions in the PFC, and premotor areas in the frontal cortex (21). We propose that the vlPFC node may have two main functions in the SN. First, the vlPFC may predict possible outcomes associated with salient stimuli identified by the AI. One example of stimulus-outcome predictions is the estimation of reward probabilities. Excitotoxic lesions in NHP area 47/12 (including its caudal portion) of macaques impaired the learning of reward probabilities after cue presentation (18). Neurons in a similar location (mid-caudal area 47/12) anticipate and predict information seeking to resolve uncertainty about future rewards and punishments (20). Caudal 47/12 shows high activation during stimulus-outcome updating when varying the visuospatial cues (19). Second, the vlPFC may be responsible for preparing appropriate behavioral responses later selected by the dACC. Two experiments in NHP show the involvement of the caudal vlPFC in this process. During win-stay/lose-shift tasks, voxels in the macaque caudal 47/12 show high activation while encoding appropriate decisions (63). In marmosets, excitotoxic lesions in area 47/12 (including its caudal portion) reduced coping mechanisms to salient negative stimuli (e.g., a fake predator in the experimental environment) (64, 65). Moreover, caudal 47/12 connects with other portions of the vlPFC associated with goal-directed movements (50, 66), which may facilitate the planning of appropriate motor responses.

Studies in humans show that AI is specifically responsible for stimulus processing, and the vlPFC is associated with stimulus-outcome predictions and response preparation. For example, a meta-analysis of stop-signal tasks (SSTs) identified independent activation clusters within the AI and vlPFC (67). The authors then trained an independent cohort undergoing a new SST and evaluated the fMRI activity in these two clusters. The AI was associated with the identification of salient information (unsuccessful trials), and the vlPFC was responsible for response implementation (inhibitory behaviors) (67). In a similar experiment, the same research group compared auditory and visual SSTs. Consistent with the first report, the AI was responsive to cue processing while the vlPFC showed a higher role in inhibitory anticipation and implementation (68). Clinical research also supports the proposed roles of the vlPFC in the SN. Smokers present abnormal activation in area 47/12 in response to cigarette cues (69-72). Similar cue-response in the right vlPFC was also reported in gamblers (69, 70) and patients with eating disorders (73). For all these patients, the poor stimulus-outcome estimation may impair response planning (74). Consequently, they engage in habitual behaviors instead of adequate responses. Addictive behaviors are also related to impaired SN function (74). Importantly, altered functional connectivity between the dACC and FIC SN nodes in these patients is correlated with abnormal vlPFC cue-response (75). Altogether, these clinical data provide additional support in favor of the vlPFC functional relevance in the SN.

### The central role of the vlPFC in attention networks

Here we demonstrated that the caudal 47/12 is an independent node of the SN. In addition to the SN, different subregions of the vlPFC are also physiologically (7-16) and anatomically (21, 50, 51, 76, 77) associated with the main nodes of the VAN (mid and caudal area 47/12) and DAN (areas 44 and 45). Importantly, the caudal area 47/12 (SN) is highly interconnected with other portions of area 47/12 (VAN) and both areas 44 and 45 (DAN) (21, 50, 51, 76, 77) Thus, the three attention networks interface extensively within a vlPFC micro-network. This central role is augmented by the fact that the vlPFC receives input from other areas of the FC. For example, the OFC is tightly linked to the vlPFC (21, 29) and provides relevant information regarding value updating (18, 78). The vlPFC is also closely connected to the dlPFC (21, 79, 80), supporting executive control functions (3). Based on its connectivity profile, we propose the vlPFC as an integrative hub combining high cognitive processing regarding attentional stimuli and switching between the main attention networks. Specifically, the vlPFC may be the area responsible for bridging the gap between the detection (VAN) and selection (DAN) of relevant stimuli, predicting outcomes, and preparing adequate behavioral responses later coordinated by the SN. These processes together, explain the critical of the vlPFC for cognitive and behavioral flexibility (23-25).

## Methods

### Injection sites

Ten adult male macaque monkeys (eight *Macaca mulatta*, one *Macaca fascicularis*, and one *Macaca nemestrina*) were used for these tracing studies. All experiments and animal care were approved by the University Committee on Animal Resources at University of Rochester and conducted following the National Guide for the Care and Use of Laboratory Animals. Retrograde tracers were injected into the right vlPFC (Supp. Fig. 1A), including one in area 47/12, one in area 47/12o and three in area 47/12l, two in area 45, one in area 44. Surgical and histological procedures were conducted as previously described (81-84). Anterograde tracers were injected into the dACC (two injections, Supp. Fig. 1B) and the FIC (two injections, Supp. Fig. 1C). Stereotaxic coordinates for the injection sites were located using pre-surgery structural MR images. Monkeys received injections of one or more of the following bidirectional tracers: Lucifer Yellow (LY), Fluororuby (FR), or Fluorescein (FS). All tracers were conjugated to dextran amine (Invitrogen) and had similar transport properties (85).

Twelve to fourteen days after the surgery, monkeys were deeply anesthetized and perfused with saline, followed by a 4% paraformaldehyde/1.5% sucrose solution. Brains were post-fixed overnight and cryoprotected in increasing gradients of sucrose (81). Serial sections of 50 mm were cut on a freezing microtome, and one in every eight free-floating sections was processed to visualize LY, FR and FS tracers, as previously described (82-84). Sections were mounted onto gel-coated slides, dehydrated, defatted in xylene overnight, and cover slipped with Permount. In cases in which more than one tracer was injected into a single animal, adjacent sections were processed for each antibody reaction.

### Anatomical tracing analysis

We first divided the FC in 23 areas and the IC in 4 areas based on the atlas by (86), in conjunction with detailed anatomical descriptions (87-90). The rationale for using the atlas of Paxinos, Huang and Toga

(86) is the homologous labeling of regions in the macaque and human brains (91, 92). Then, FC and IC areas were grouped according to common cytoarchitectonic characteristics: area 10 (including subdivisions 10, 10d, 10l and 10m), 25, 14 (14o and 14m), 11 (11, 11m, and 11l), 13 (13, 13a, 13m, and 13l), 24 (24a, 24b, and 24c), 32, 46 (46v, and 46d), 9 (9l, 9m, 9/32, 9/46, 9/46v, and 9/46d), 8 (8/32, 8a, 8ad, 8av, and 8b), 6m (6/32, and 6m), 6d (6dc/F2, and 6dr/F7), 6v (6vc/F4, 6vr/F5, and ProM), OPAl, OPro, AI, DI, GI, and IPro.

#### Retrograde analysis

To evaluate the strength of afferent projections from the FC and IC to the vlPFC, light field microscopy under 20x objective was used to identify retrogradely labeled cells, as previously described (84, 93, 94). StereoInvestigator software (MicroBrightField Bioscience, U.S.A) was used to stereologically count cells in one of every 24 sections (1.2 mm interval). Cell counts were obtained in 19 FC/IC areas previously listed. For each case, the connectivity strength (CS) between each area and the injection site was estimated by a percent score (84):

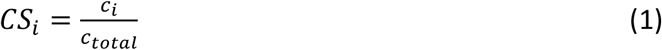

where *CS*_*i*_ is the connectivity strength for the i-th area, ci is the cell count in the i-th area, and ctotal is the total number of labeled cells across all FC/IC areas.

We also performed a random sampling analysis to evaluate the connectivity strengths expected by chance in each area. For this, the total number of cells in each case was randomly assigned to each FC or IC area with a probability given by the volume of the area. The connectivity strength was then calculated according to Equation 1. This procedure was repeated 106 times to create a random distribution. The 95% confidence intervals (CI) of these random distributions were computed for each one of the 19 FC/IC areas in each case.

Finally, we calculated the Spearman correlation between the connectivity strength across the 19 FC/IC areas in both caudal vlPFC cases.

#### Anterograde analysis

For the dACC and FIC injection cases, dark field light microscopy under 1.6x, 4x, and 10x objectives was used with Neurolucida software (MicroBrightField) to trace outlines of dense or light focal projections to the caudal vlPFC. “Dense projections” were characterized by condensed groups of fibers visible at 1.6x with discernible boundaries (93, 94). Condensed group of fibers where individual terminals could be discerned were labeled as “light projections” (Fig. 2B, bottom).

### Functional neuroimaging

#### Macaque dataset

The macaque fcMRI maps were generated from one adult monkey (*Macaca mulatta*, female, age 7 years, weight 6.4 kg; NKI dataset) from the PRIMatE Data Exchange (PRIME-DE) consortium (39). This monkey has four anesthetized scanning sessions with monocrystalline iron oxide ferumoxytol (MION) as the contrast agent. Each session consists of 4–8 scans (8-min per scan). The NKI Institutional Animal Care and Use Committee (IACUC) protocol approved all methods and procedures.

#### Macaque data acquisition

All MRI data were collected using an 8-channel surface coil adapted for monkey head scanning on a 3.0 Tesla Siemens Tim Trio scanner (Siemens, Erlangen, Germany). Structural images were obtained using a T1-weighted sequence (TR=2500 ms, TI=1200 ms, TE=3.87 ms, FA= 8°, 0.5 × 0.5 × 0.5-mm voxels). Functional data were collected using a gradient-echo EPI sequence (TR=2000 ms, TE=16.6 ms, FA= 45°, 1.5 × 1.5 × 2-mm voxels, 32 slices, FOV=96 × 96 mm). Monocrystalline iron oxide ferumoxytol (MION) solution was injected at iron doses of 10 mg/kg IV before the MRI scanning. The monkey was sedated with an initial dose of atropine (0.05 mg/kg IM), dexdomitor (0.02 mg/kg IM), and ketamine (8 mg/kg IM) intubated and maintained at 0.75% isoflurane anesthesia during the scanning.

#### Macaque data preprocessing

Structural data preprocessing included the following steps: (1) spatial noise removing and bias field correction using ANTs, (2) brain extraction and segmentation into gray matter, white matter and cerebrospinal fluid using FSL and FreeSurfer, and (3) reconstructing the native white matter and pial surface using FreeSurfer.

Functional data were preprocessed by the following steps: (1) first 5 frames of BOLD data were dropped, constant offset and linear trend over each run were removed, (2) six parameters were obtained by motion correction with a rigid body registration algorithm, (3) spatial smoothing was performed with a Gaussian kernel of FWHM 2mm, (4) each run was then normalized for global mean signal intensity, (5) a band-pass temporal filter was applied to retain frequencies to 0.01 Hz < 0.1 Hz, (6) head motion, whole-brain signal, ventricular and white matter signals were removed through linear regression, (7) the preprocessed fMRI data was then registered to the macaque MNI template and down-sampled to the 1mm resolution for further analysis.

#### Macaque functional connectivity analysis

Seven 3 mm radius seeds were placed in corresponding locations to our vlPFC injection sites according to the macaque MNI space (Fig. 3A, Supp. Table 1) (95). The resulting connectivity matrices were later linear projected to the macaque MNI space with 0.25 mm resolution. We created a mask for the dACC, and FIC based on the cluster of cells identified using the retrograde tract-tracing. Then, we used the Fisher r-to-s transformation to correct the correlation values of each voxel, and the functional connectivity strength between each mask and a seed was calculated as the absolute average value within the respective mask.

To evaluate if our results were different from the chance level, we created a random distribution of connectivity strengths in each mask. For this, we performed a random permutation of voxels across the brain volume. Then, we calculated the functional connectivity strength between each mask and seed as previously described. This procedure was repeated 10^6^ times to create a random distribution. The 95% confidence intervals (CI) of these random distributions were computed for each mask in each case.

As a secondary analysis, we placed five 3 mm radius seeds inside and outside the dACC mask, according to the macaque MNI space (Fig. 3B, Supp. Table 1) (95). Then, we calculated the functional connectivity between each vlPFC and dACC seeds. Before analysis, these values were also r-to-z transformed.

#### Human dataset

For the cross-species functional connectivity analysis, we used a dataset consist of 1000 young, healthy adult participants (mean age 21.3 ± 3.1 years; 427 males) from the Brain Genomics Superstruct Project (GSP) (40). Each participant performed one structural MRI run and 1-2 resting-state fMRI runs (6 min 12 s per run). All participants provided written informed consent following guidelines set by the Institutional Review Boards of Harvard University or Partners Healthcare.

#### Human data acquisition

All MRI data were acquired using a 12-channel head coil on matched 3T Tim Trio scanners (Siemens, Erlangen, Germany). Structural data were obtained by a multi-echo T1 weighted gradient-echo image sequence (TR = 2200 ms, TI = 1000 ms, TE = 1.54 ms for image 1 to 7.01 ms for image 4, FA = 7°, 1.2 × 1.2 × 1.2-mm voxels, and FOV = 230). Resting-state functional MRI images were collected using the gradient-echo EPI sequence (TR = 3000 ms, TE = 30 ms, flip angle = 85°, 3 × 3 × 3-mm voxels, FOV = 216, and 47 axial slices collected with interleaved acquisition). Participants were instructed to stay awake and keep their eyes open during the scanning.

#### Human data preprocessing

Structural MRI data were preprocessed using the “recon-all” pipeline from FreeSurfer software package. The individual surface mesh was reconstructed and registered to a common spherical coordinate template.

Functional MRI data were processed using a previously described preprocess pipeline, including: (1) slice timing correction using SPM, (2) head motion correction by FSL, (3) normalization for global mean signal intensity across runs, (4) band-pass filtering (0.01-0.08 Hz), and (5) regression of motion parameters, whole-brain signal, white matter signal, and ventricular signal. The preprocessed fMRI data were then registered to the MNI152 template and downsampled to a 2 mm spatial resolution. Spatial smoothing with a 6 mm FWHM kernel was performed on the fMRI data within the brain mask.

#### Human functional connectivity analysis

Eleven 5 mm radius seeds were placed in corresponding locations to our vlPFC injection sites according to the MNI152 template (Fig. 4A, Supp. Table 2). After the creation of a dACC and an FIC mask in homologous positions of the cell clusters identified in the macaque retrograde data, correlation values were r-to-z transformed, and the connectivity strength between each seed and mask was calculated as the absolute average value within the respective mask. The same random permutation approach used for the monkey data was repeated here.

As a secondary analysis, we placed seven 5 mm radius seeds inside and outside the dACC mask, according to the macaque MNI152 template (Fig. 4B, Supp. Table 2) and calculated the seed-to-seed connectivity between the vlPFC and dACC. The Fisher r-to-z transformation was also applied to these results. Finally, we repeated both analyses using 3 mm radius seeds to ensure that our results were not driven by the seed size (Supp. Fig. 5).

