## Supplementary Material for "Anatomical and functional connectivity support the existence of a salience network node within the caudal ventrolateral prefrontal cortex"

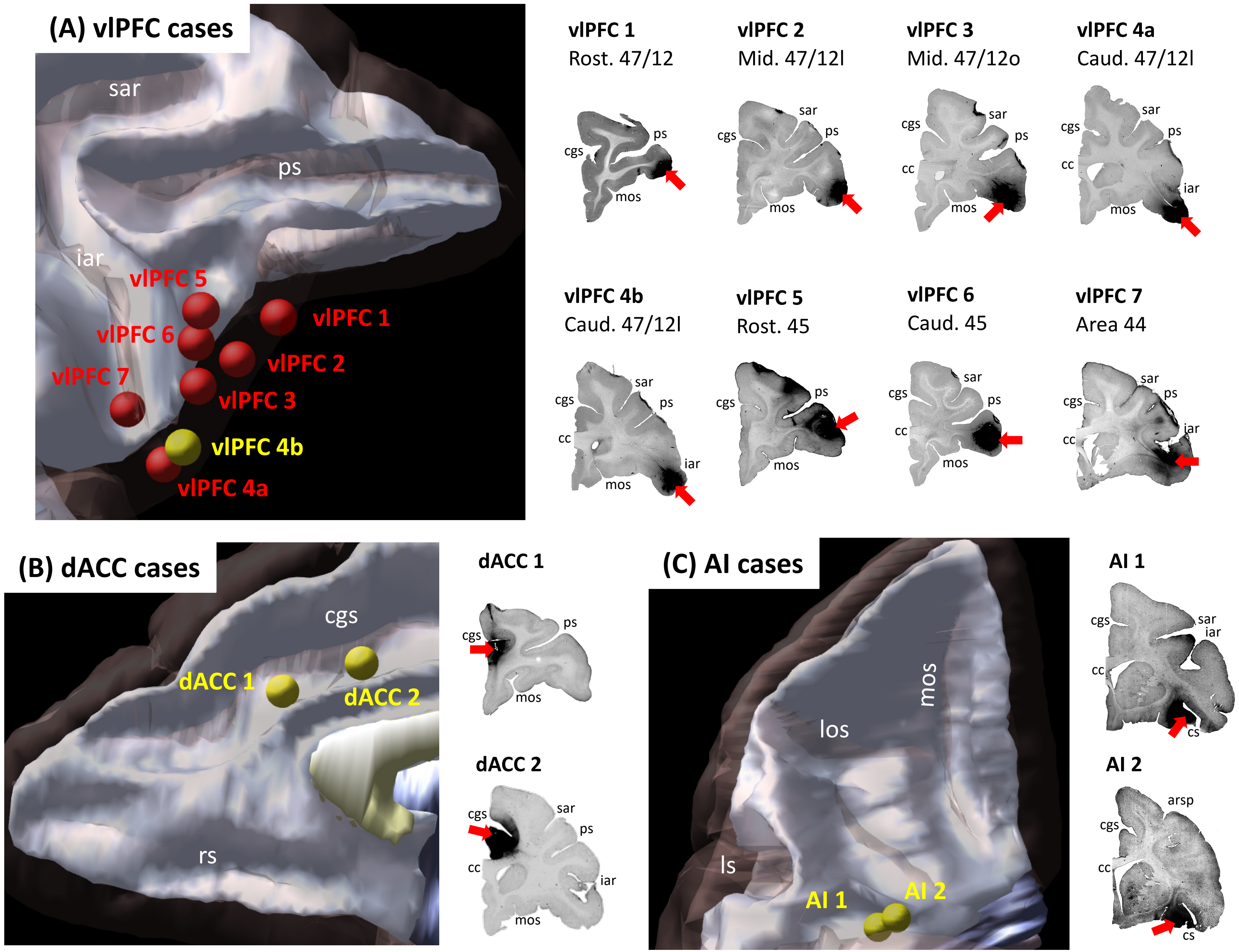


**Supp. Fig. 1 - Injection sites. (A)** Location of 8 injection locations in the vlPFC selected for retrograde analysis. Seven cases were analyzed as the main results (red), and one case was used as validation (yellow). Injection locations in **(B)** the dACC and **(C)** the AI selected for anterograde validation of the salience node.
*Abbreviations:* arsp = arcuate sulcus spur; cc = corpus callosum; cgs = cingulate sulcus; cs = circular sulcus; iar = inferior arcuate sulcus; ls = lateral sulcus; los = lateral orbital sulcus; mos = medial orbital sulcus; ps = principal sulcus; rs = rostral sulcus; sar = superior arcuate sulcus.

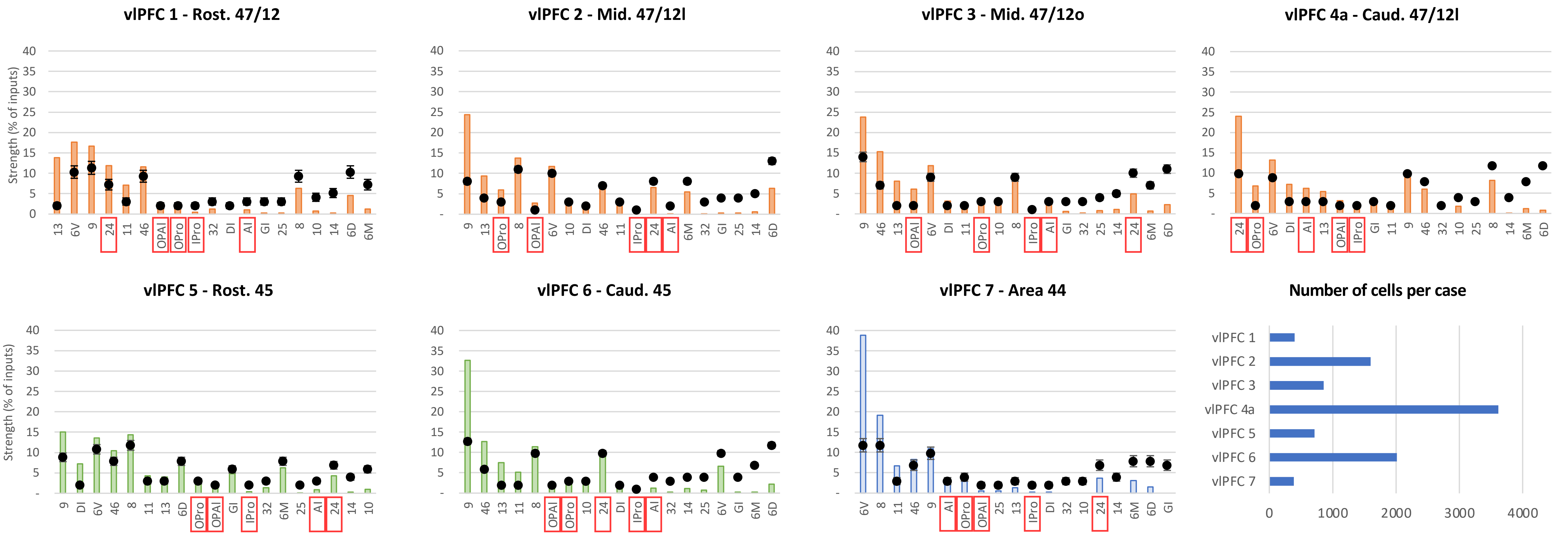


**Supp. Fig. 2 – Strength of projections from the frontal and insular cortices to different regions of the vlPFC.** Bars are sorted from stronger to weaker projections in each case. Orange bars illustrate cases with injections in area 47/12, green bars in area 45, and blue bars in area 44. Black dots show the average and standard-deviation of random sampling from the respective areas in each case. Red squares highlight cytoarchitectonic areas relevant for the Salience Network.

**(A) projections from the frontal and insular cortices to different the rostral area 47/12**

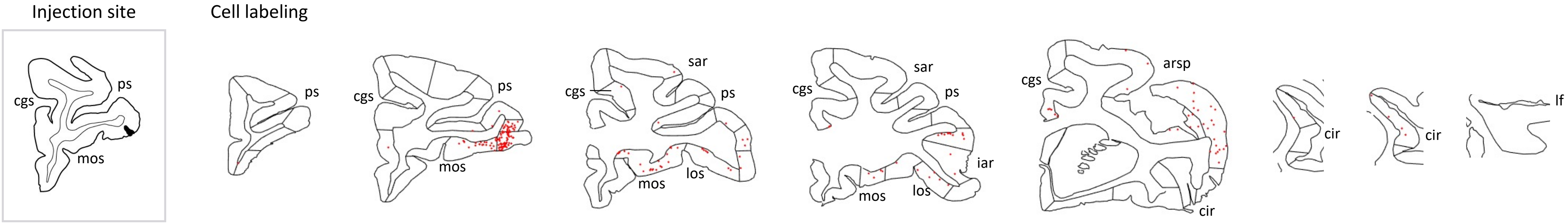


**(B) projections from the frontal and insular cortices to different the mid area 47/12l**

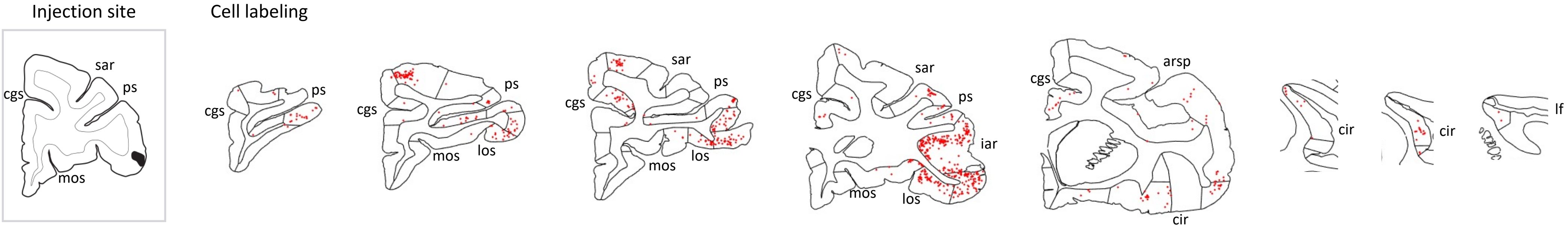


**(C) projections from the frontal and insular cortices to different the mid area 47/12o**

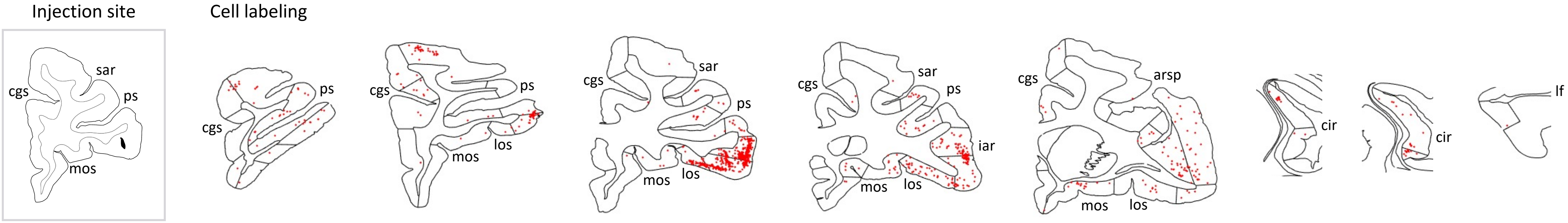


**(D) projections from the frontal and insular cortices to different the caudal area 47/12l**

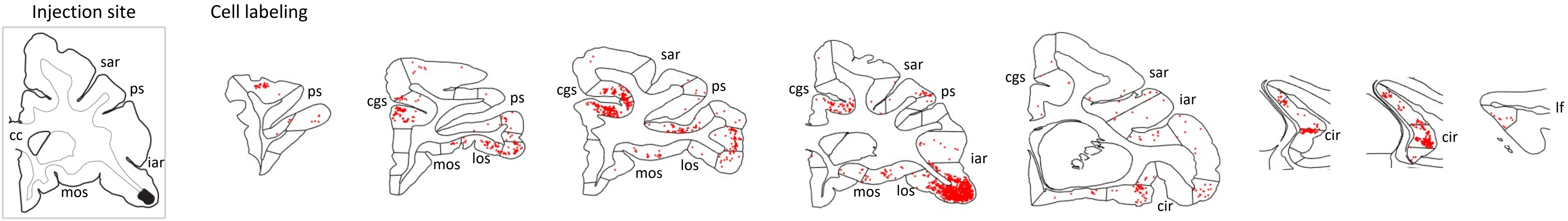

**(E) projections from the frontal and insular cortices to different the rostral area 45**

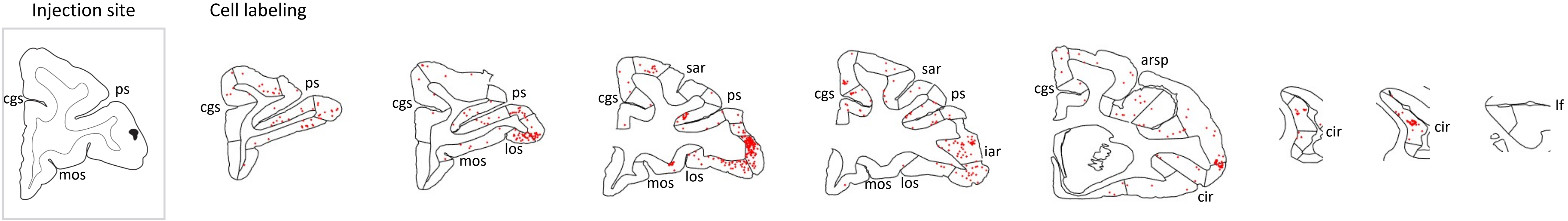


**(F) projections from the frontal and insular cortices to different the caudal area 45**

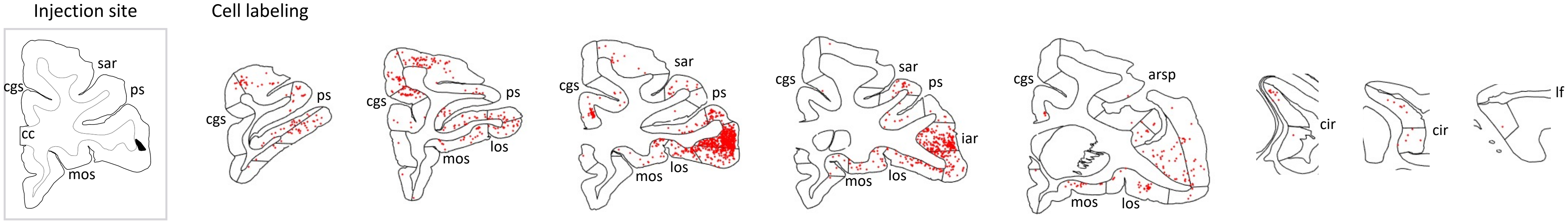


**(G) projections from the frontal and insular cortices to different the area 44**

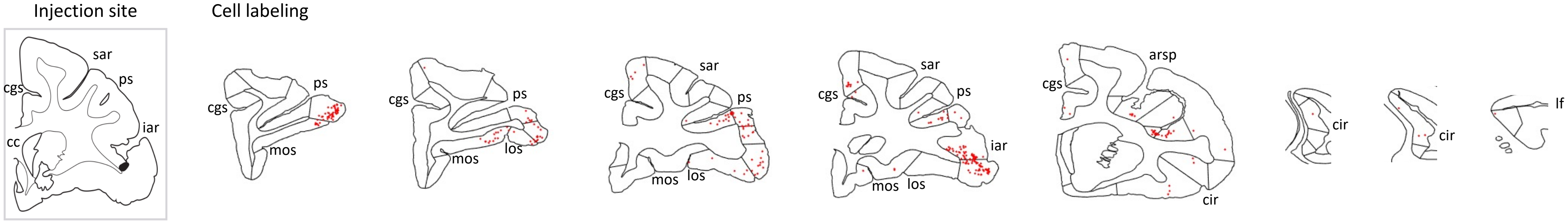


**Supp. Fig. 3 – (A-G) Injection locations and retrogradely labeled cells in the main vlPFC cases.** Each line contains rostral to caudal coronal sections and the respective labelled cells (red dots) from one case. Sections of the same column have matching locations along the rostro-caudal axis. The last three columns correspond to the IC.

*Abbreviations:* arsp = arcuate sulcus spur; cgs = cingulate sulcus; cir = circular sulcus; iar = inferior arcuate sulcus; lf = lateral fissure; los = lateral orbital sulcus; mos = medial orbital sulcus; ps = principal sulcus; sar = superior arcuate sulcus.

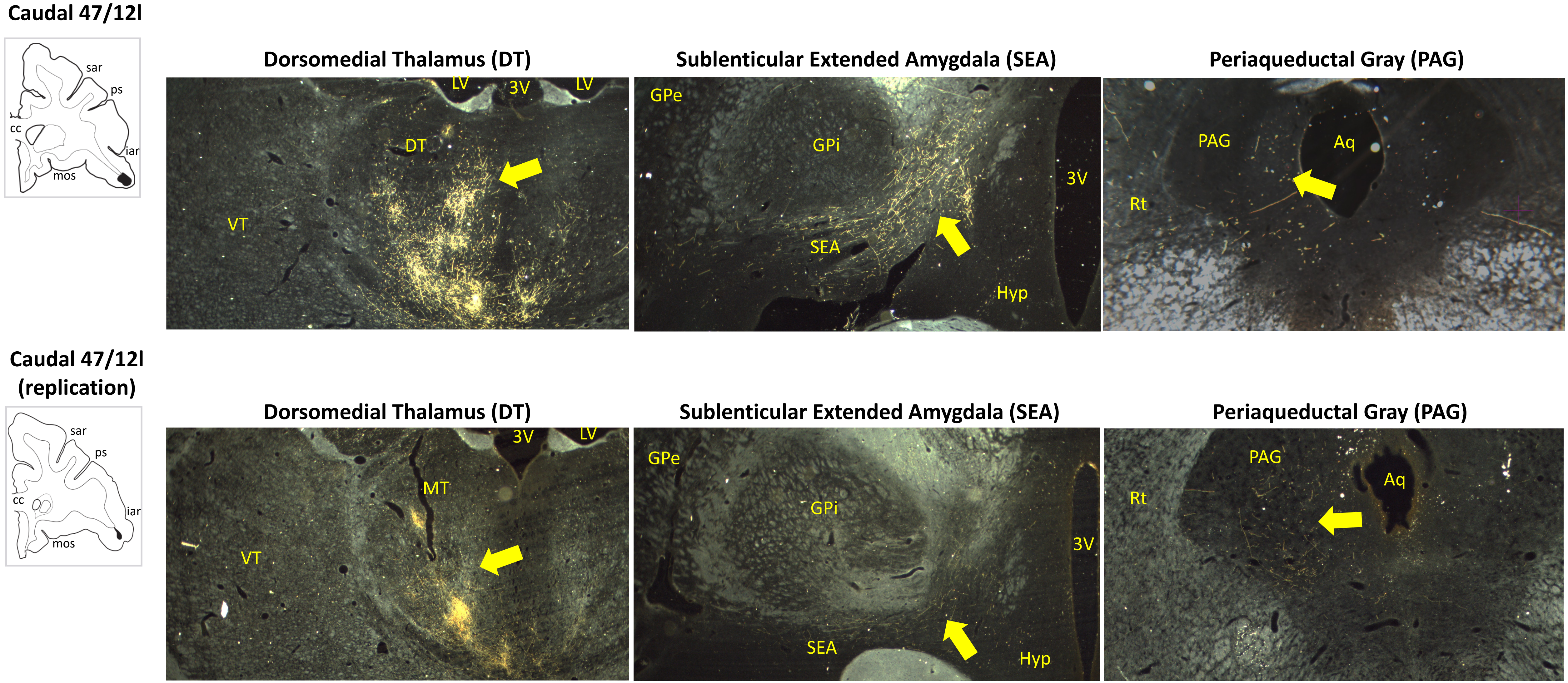


**Supp. Fig. 4 – Terminal fields from injections in caudal vlPFC area 47/12 within the dorsomedial thalamus (1x amplification), sublenticular extended amygdala (1x), and periaqueductal gray (2x).**

*Abbreviations:* 3V = 3rd ventricle; Aq = Aqueduct; DT = Dorsomedial thalamus; GPe = Globus pallidus ext.; GPi = Globus pallidus int.; Hyp = Hypothalamus; LV = lateral ventricle; PAG = Periaqueductal gray; Rt = Reticular formation; SEA = Sublenticular extended amygdala; SP = superior peduncle; VT = Ventralposterior thalamus.

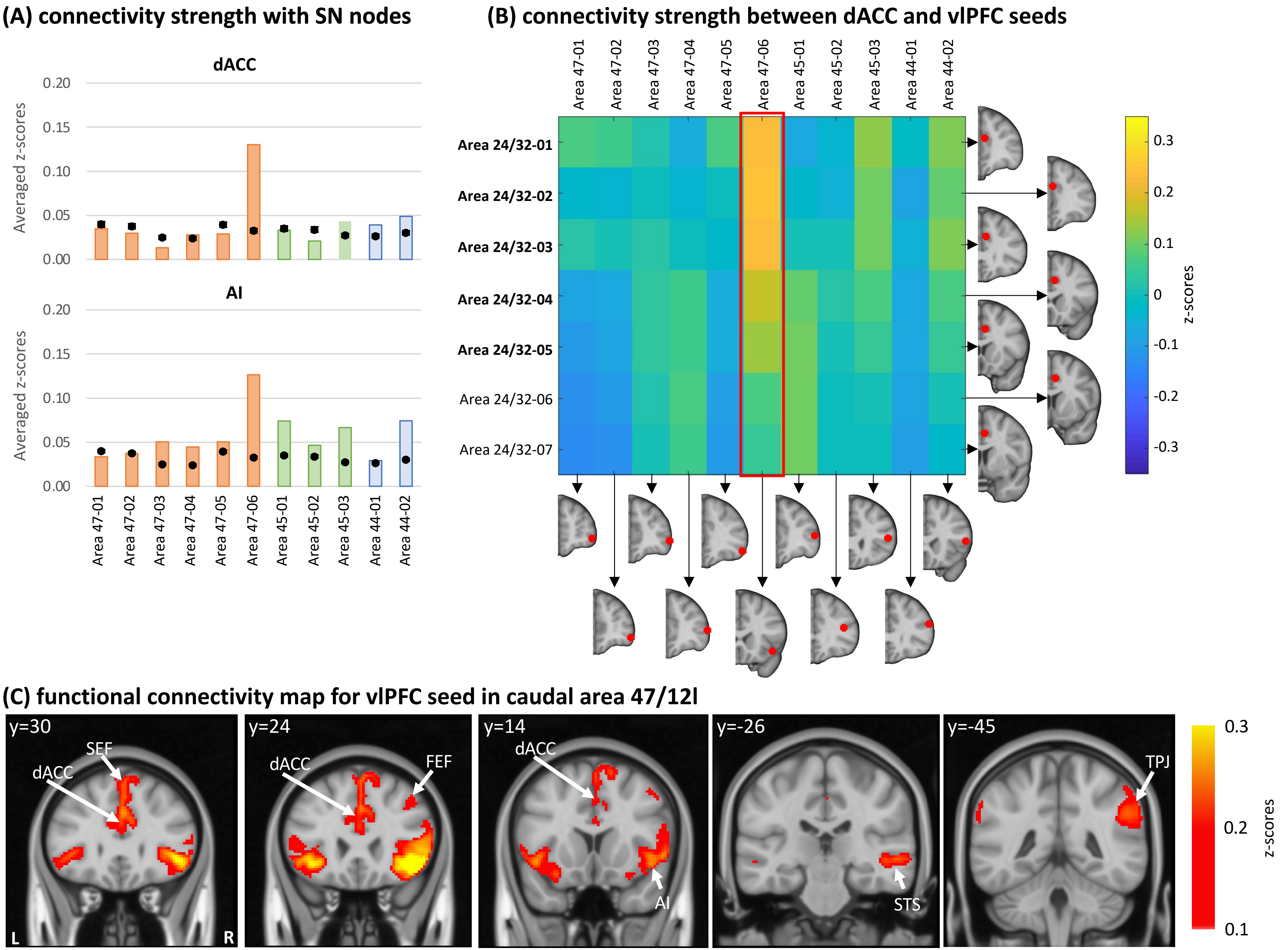


**Supp. Fig. 5 – Human fMRI analysis using a seed radius of 3 mm. (A)** Average connectivity strength (z-scores) between vlPFC seeds and the dACC and AI masks. Orange bars illustrate cases with injections in area 47/12, green bars in area 45, and blue bars in area 44. Black dots show the average and standard-deviation of the voxel permutation analysis. **(B)** Connectivity strength (z-scores) between dACC and vlPFC seeds. In bold the seeds overlapping with the dACC mask. **(C)** Different views of the voxel distribution for the caudal 47/12l seed. All coordinates

**Macaque MNI coordinates**

| Seed | x | y | z |
|---|---|---|---|
| | ***vlPFC seeds*** | | |
| *Rost. 47/12* | 14.50 | 19.00 | 6.00 |
| *Mid. 47/12l* | 18.50 | 13.50 | 1.00 |
| *Mid 47/12o* | 16.00 | 13.25 | 0.00 |
| *Caud/ 47/12l* | 18.75 | 11.00 | -3.50 |
| *Rost. 45* | 17.75 | 13.50 | 4.25 |
| *Caud. 45* | 18.75 | 12.25 | 2.25 |
| *Area 44* | 20.00 | 9.75 | 0.75 |
| | ***dACC seeds*** | | |
| *Area 24* | 2.00 | 17.50 | 8.50 |
| *Area 24* | 4.00 | 14.50 | 10.00 |
| *Area 24* | 4.00 | 12.00 | 10.50 |
| *Area 24* | 1.00 | 10.00 | 10.50 |
| *Area 24* | 1.00 | 7.00 | 11.50 |

**Supp. Tab. 1** – ROI centers for macaque fMRI analysis.

**Human MNI coordinates**

| Seed | x | y | z |
|---|---|---|---|
| | ***vlPFC seeds*** | | |
| *Area 47-01* | 44.00 | 54.00 | -6.00 |
| *Area 47-02* | 48.00 | 48.00 | -8.00 |
| *Area 47-03* | 52.00 | 44.00 | -2.00 |
| *Area 47-04* | 54.00 | 40.00 | 6.00 |
| *Area 47-05* | 52.00 | 34.00 | -10.00 |
| *Area 47-06* | 44.00 | 22.00 | -10.00 |
| *Area 45-01* | 50.00 | 38.00 | 8.00 |
| *Area 45-02* | 40.00 | 32.00 | 16.00 |
| *Area 45-03* | 50.00 | 26.00 | 12.00 |
| *Area 44-01* | 54.00 | 30.00 | 22.00 |
| *Area 44-02* | 54.00 | 22.00 | 10.00 |
| | ***dACC seeds*** | | |
| *Area 24/32* | 6.00 | 40.00 | 20.00 |
| *Area 24/32* | 2.00 | 34.00 | 26.00 |
| *Area 24/32* | 6.00 | 30.00 | 30.00 |
| *Area 24/32* | 6.00 | 20.00 | 34.00 |
| *Area 24/32* | 6.00 | 16.00 | 36.00 |
| *Area 24/32* | 6.00 | 10.00 | 40.00 |
| *Area 24/32* | 4.00 | 6.00 | 42.00 |

**Supp. Tab. 2** – ROI centers for human fMRI analysis.
